# Ultrasensitive in vivo infrared spectroscopic imaging via oblique photothermal microscopy

**DOI:** 10.1101/2024.10.02.616360

**Authors:** Mingsheng Li, Sheng Xiao, Hongli Ni, Guangrui Ding, Yuhao Yuan, Carolyn Marar, Jerome Mertz, Ji-Xin Cheng

## Abstract

In vivo IR spectroscopy faces challenges due to poor sensitivity in reflection mode and low resolution at micrometer scale. To break this barrier, we report an oblique photothermal microscope (OPTM) to enable ultrasensitive IR spectroscopic imaging of live subjects at sub-micron resolution. Classic photothermal measurement captures only a small fraction of probe photons through a pinhole to extract the photothermal signal. Instead, OPTM uses a differential split detector placed on the sample surface to collect 500-fold more photons and suppress the laser noise by 12 fold via balanced detection. Leveraging its improved sensitivity, OPTM enables low-dose IR imaging of skin without photodamage. Depth-resolved in vivo OPTM imaging of metabolic markers beneath mouse and human skin is shown. Furthermore, we demonstrate in vivo OPTM tracking of topical drug contents within mouse and human skin. Collectively, OPTM presents a highly sensitive imaging platform for in vivo and in situ molecular analysis.

## Introduction

Label-free imaging of chemicals in live animals and human subjects is crucial for both basic research and clinical translation. Vibrational spectroscopic imaging has emerged as a highly sensitive, label-free platform to visualize molecular contents in living systems, advancing the study of biology and medicine (*1*). In particular, spatially offset Raman spectroscopy and coherent Raman scattering microscopy have been developed to image chemicals in the skin layers of live subjects (*2-4*). Compared to spontaneous Raman scattering, infrared (IR) absorption provides a roughly 8-order-of-magnitude larger cross section, offering a more sensitive method for imaging chemicals using their fingerprint signals (*5*). Since William Coblentz introduced the IR spectrometer in the early 1900s to explore molecular structures (*6*), IR spectroscopy has been significantly advanced. Among the various techniques, Fourier-transform infrared (FTIR) spectroscopy is extensively used in the fields of biological research, function materials, and pharmaceuticals (*7, 8*). The invention of quantum cascade laser as a room-temperature semiconductor laser has facilitated highly sensitive IR spectroscopic imaging (*9, 10*). Traditional IR spectroscopic imaging measures the loss of IR photons, which is not suitable for imaging live animals because IR photons are attenuated and cannot penetrate through or reflected by an intact animal body. Photoacoustic IR microscopy circumvents the issue of low photon collection efficiency by using the weak-scattering photoacoustic wave as a readout of IR absorption (*11-13*). However, because IR photons attenuate in the acoustic coupling medium, photoacoustic IR microscopy cannot operate in an epi-mode, making it not applicable for universal in vivo imaging scenarios. FTIR and photoacoustic IR also face limitations in spatial resolution due to the diffraction limit of long IR wavelengths, making them inefficient for nanoscale chemical visualization in vivo. Using a near-field probe strategy to overcome the IR diffraction limit, atomic force microscope-infrared (AFM-IR) spectroscopy achieves 10-nm resolution (*14-16*). However, AFM-IR’s limited penetration depth makes it unsuitable for volumetric imaging of biological samples.

Recently developed mid-infrared photothermal (MIP) microscopy addresses these limitations by using a visible light to probe the mid-infrared-induced thermal effects (*17-19*). Over the past few years, advances have been made towards widefield measurement (*20-23*), video-rate imaging speed (*24, 25*), micromolar-level sensitivity in fingerprint and silent windows (*17, 26, 27*), and 3D tomographic imaging capability (*28-30*). Despite these advances, in vivo MIP imaging has not yet been feasible due to limited sensitivity in visualizing chemical content in live animals. The epi-detection geometry is necessary for in vivo imaging scenarios because photons cannot penetrate through the animal body. In a classic epi-detected MIP microscope (**Figure 1a**), the pump and probe beam are coaligned and focused by a reflective objective onto a sample (*31*). The scattered photons from the sample are collected by the same objective. An iris is employed before a remote photodetector to maximize the photothermal signals. However, the probe photons experience enormous scattering events in complex tissues, resulting in scattered photons losing their original propagation directions. Thus, using an objective and an iris to collect scattered photons is ineffective. While classic MIP can visualize chemical content in opaque samples, such as pharmaceutical tablets (*31, 32*) and thin tissue slices (*33*), it is not sensitive for in vivo measurement due to significant loss of highly scattered probe photons during back-propagation to the remote detector.

**Figure 1.**
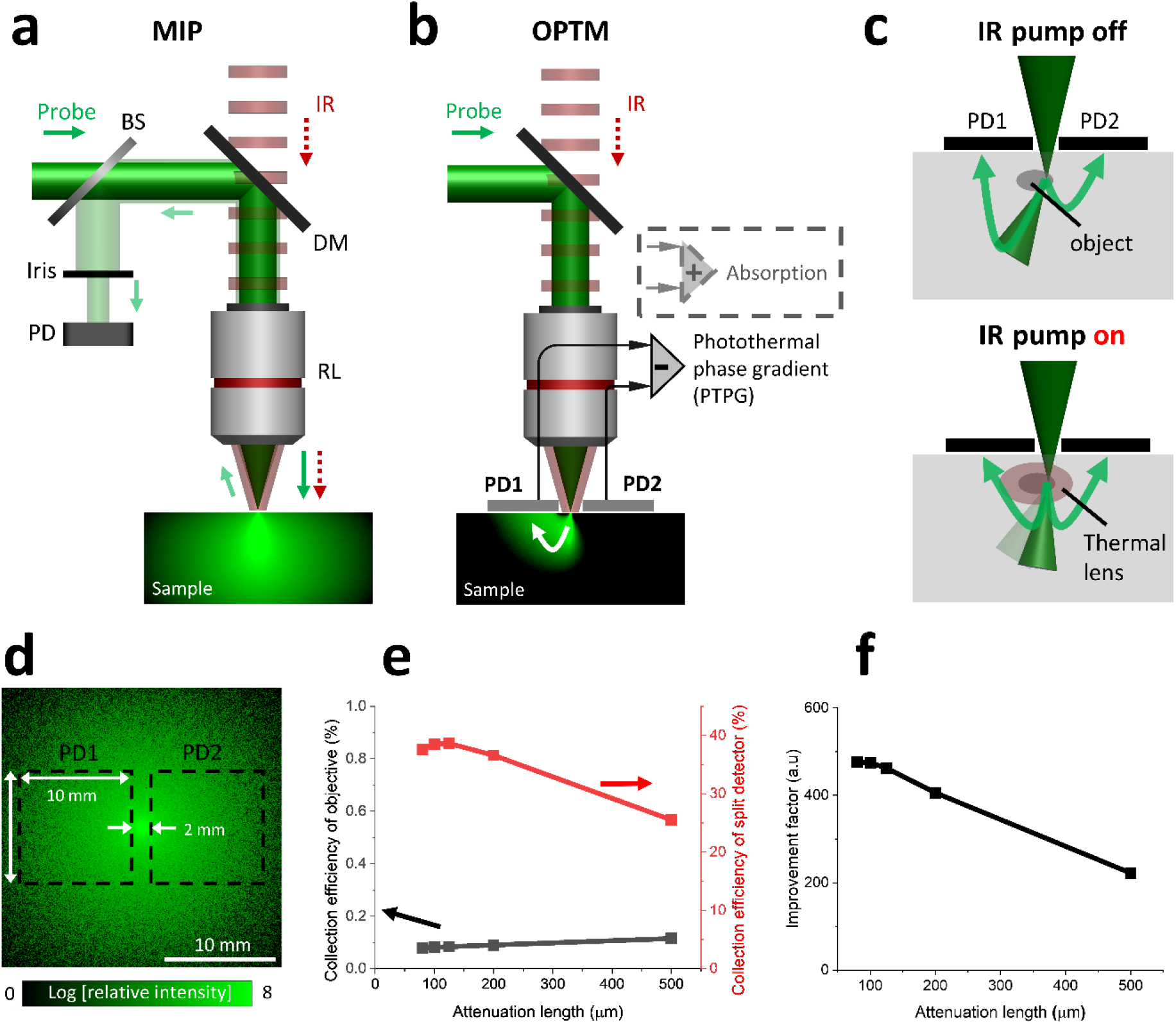
OPTM principle and simulation results. **a**, schematic of mid-infrared photothermal (MIP) microscope. **b**, schematic of oblique photothermal microscope (OPTM). Photons propagating to PD1 are shown in the sample box. **c**, the principle of OPTM, illustrating the probe propagations at the status of IR on and off. **d**, Monte Carlo simulation result of photon distribution on the surface of a uniform scattering layer. **e**, photon collection efficiency of split detector and objective. **f**, the improvement factor in photon collection efficiency with oblique detection compared to the classic method. BS: beam splitter. DM: dichroic mirror. PD: photodiode. RL: reflective objective.

Here, we report an oblique photothermal microscope (OPTM) to enable high-sensitivity in vivo infrared spectroscopic imaging at sub-micron resolution (**Figure 1b**). Instead of an iris before a remote detector, oblique photothermal detection is achieved by placing a differential split detector above the sample surface to collect epi-propagated probe photons with high efficacy. After multiple scattering within tissue, the forward scattered photons from the focus are redirected to the backward direction before reaching to the split detector. The propagation path of collected photons aligns obliquely from the focus to the split detector. Such oblique optical detection or illumination using two fibers provides phase gradient information of samples for improved contrast, in which phase gradient signal is proportional to refractive index variations (*dn*⁄*dx, n* is refractive index of sample, *x* is the lateral distance) (*34-37*). Yet, oblique detection itself only gives the phase gradient information, without molecular sensitivity.

In OPTM, photothermal expansion of objects decreases its refractive index, thus reducing the deflection angle of the original probe path. Consequently, with photothermal effect, one photodiode receives more photons while the other receives fewer. Thus, photothermal modulations from each half of the split detector have opposite signs. Subtracting the two signals of the split detector not only enhances the photothermal signal but also suppresses the laser noise via a balanced detection operation. In comparison, the classic photothermal microscopy only collects a small fraction of back scattered photons from the focus (*31*), which could be overwhelmed by the laser noise and detector noise. As shown below, our method increases the photon collection efficiency by 500 times and suppresses the laser noise by a factor of 12 via balanced detection. Leveraging its enhanced sensitivity, OPTM allows low-dose IR spectroscopic imaging of animal skin, thereby avoiding the risk of photodamage. Consequently, OPTM enables in vivo IR spectroscopic imaging of metabolic markers within animal and human skin. Moreover, OPTM allows for depth-resolved monitoring of topical drugs under mouse and human skin, unveiling the topical drug pathways and allowing quantitative evaluation of drug delivery efficiency. These advances highlight OPTM’s potential in biomedical research and in situ molecular analysis.

## Results

### OPTM principle and simulation results

In an epi-detected MIP microscope (*31*), IR and visible probe beam are combined via a dichroic mirror, focused by a reflective objective, then delivered onto a sample (**Figure 1a**). The probe photons collected by the same objective are filtered by an iris to maximize the photothermal signals. In OPTM, both pump and probe beams are coaxially aligned. A split detector is positioned upon the sample surface, allowing it to collect more epi-propagated probe photons (**Figure 1b**). The photon propagation path aligns obliquely from the focus to the detector. The difference in intensity between the two halves of the split detector reveals the phase gradient information of an object within its surrounding medium, while the sum provides optical absorption information of the same object (*35, 38*). Upon photothermal modulation, the intensity difference between both detectors yields photothermal phase gradient (PTPG) contrast. For comparison with PTPG images, absorption images are acquired simultaneously by taking the sum of the split detector’ signals.

With the IR pump off, the object deflects the beam path due to different refractive indices between the object and its environment (**Figure 1c, upper**). The intensity difference between the split detector is proportional to the variation of refractive index in samples (*dn*⁄*dx, n* is refractive index of sample, *x* is the lateral distance) (*34-37*). With the IR pump on, the object absorbs IR photon energy, undergoes thermal expansion, and creates a thermal lens in its surrounding environment (**Figure 1c, lower**). Because the thermal lens decreases the refractive index and increases the dimension of the object, it reduces the deflection angle of the original probe path. Compared to the IR-off status, one photodiode receives more photons while the other receives fewer photons. Consequently, the photothermal signals from each half of the split detector have opposite signs. Therefore, the subtraction operation enhances the photothermal signals. Meanwhile, equivalent to balanced detection, the subtraction operation suppresses the common-mode fluctuation which is contributed by the probe laser noise (*39, 40*).

To quantitatively demonstrate an improved photon collection efficiency, we conducted a Monte Carlo simulation to analyze the propagation of four million photons within a uniform scattering layer (**Figure 1d**) (*41, 42*). On the layer surface, the objective only collects the central photons within a finite angle of 30 degree corresponding to the NA of the objective. In contrast, the split detector collects the scattered photons in a larger area as indicated with black dash box in **Figure 1d.**

We summed the photons on the detection areas of the split detector and the objective, divided it by the total photons on sample surface, and generated the photon collection efficiencies of the split detector and objective, as indicated by red and black curves in **Figure 1e.**

Although the slit within split detectors causes the leakage of photons, the collection efficiency of split detector can still reach to 39% while that of the objective is only ∼0.1%. We computed the improvement factor by dividing the collection efficiency of the split detector to that of objective. Our results indicate oblique detection offers around 500-times larger collection efficiency of scattered photons than the objective detection method (**Figure 1f**). In summary, instead of pinhole filtering, OPTM uses a split detector to collect 500-fold more epi-propagated probe photons. Harnessing the inversed photothermal modulations on the split detector, OPTM employs the phase-gradient signal as an efficient readout of photothermal contrast by subtracting the signals from the split detector.

### OPTM instrumentation and signal processing

As illustrated in **Figure 2a**, the OPTM combines a pulsed IR laser with a visible continuous wave laser as the pump and probe sources. The combined beams are delivered to the scan unit for beam scanning and focused by a reflective objective onto sample. The scan unit includes a 2D galvo scanner for scanning laser beams, followed by reflective relay optics that includes two concave mirrors (**Figure 2b**). The use of reflective relay optics and a reflective objective mitigates the achromatic aberrations of the visible and IR beams (*24, 43*). To enable oblique photothermal detection in the setup, a split detector consisting of two identical photodiodes is placed above sample surface (**Figure 2a**). A slit between the detectors allows the laser beam to pass through. To facilitate oblique photothermal detection, we engineered a detection circuit on a customized PCB (**Figure 2c-d**). Because oblique detection collects more probe photons, to avoid saturation, the split detector is biased by a 100-V DC source (V_B_) to increase the saturation threshold of photodiodes. A RC lowpass filter is used to remove the high-frequency noise in the DC source. The signals from two halves of split detector are recorded independently, following the signal processing flowchart shown below. For comparison, an epi-detected MIP microscope is shown in **Figure S1**, where a beamsplitter is used to reflect the backward-propagated photons, then filtered by an iris before reaching a remote photodiode.

**Figure 2.**
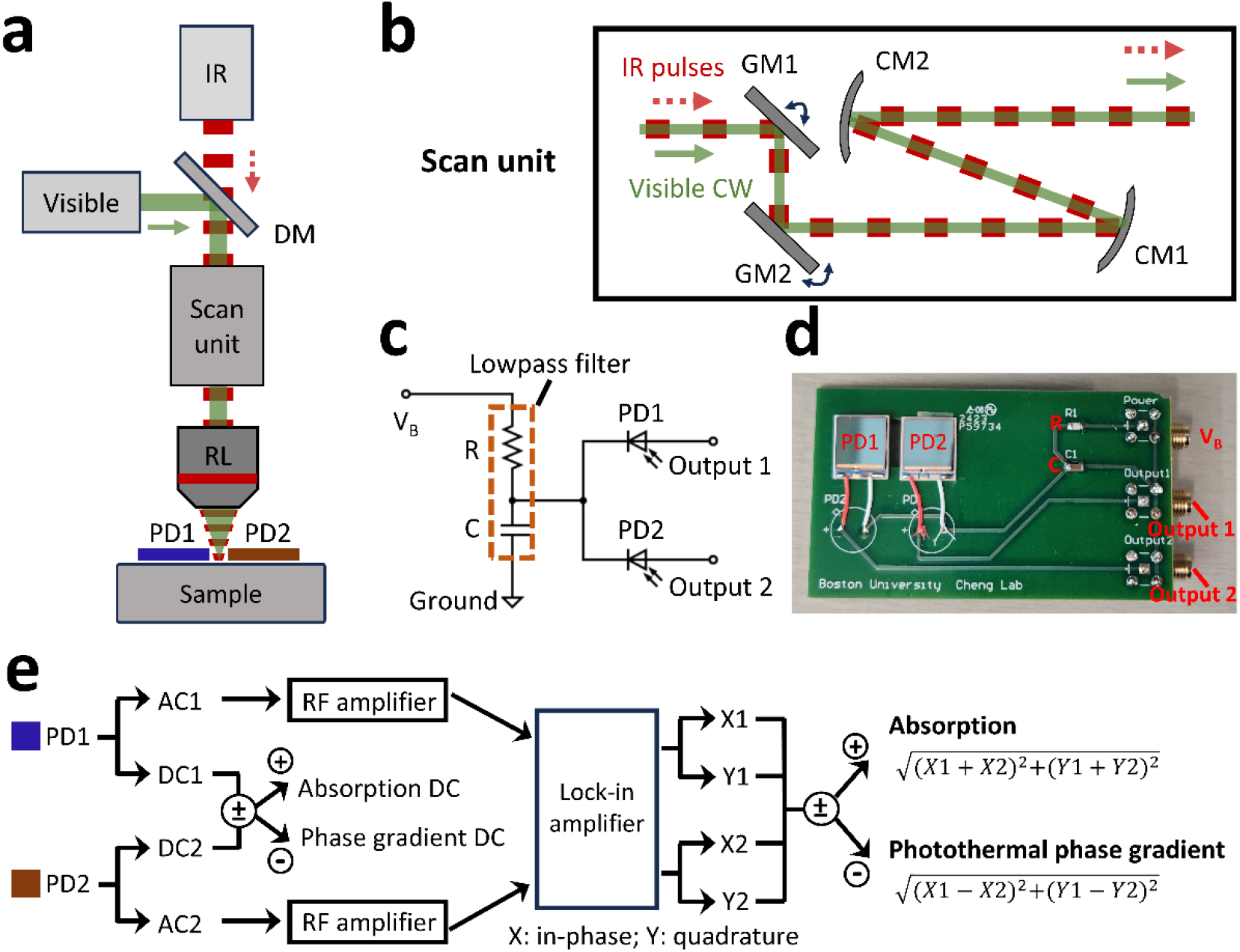
OPTM instrumentation and signal processing. **a**, an oblique photothermal microscope (OPTM). **b**, details of scan unit. **c**, circuit diagram of split detector for collecting photocurrent from each photodiode independently. **d**, photo of split detector. **e**, signal processing flowchart of generating absorption and photothermal phase gradient images from split detector’s signals. DM: dichroic mirror. RL: reflective objective. PD: photodiode. CW: continuous wave. GM: galvo mirror. CM: concave mirror.

To generate photothermal phase gradient and absorption images, we developed a signal processing flowchart as illustrated in **Figure 2e.**

Signals from each half of the split detector are split into AC and DC components. The DC components are recorded by a data acquisition card (DAQ). The sum of DC channels yields optical absorption information of probe wavelength, while their difference provides phase gradient of the objects. Each AC component is amplified and delivered to a lock-in amplifier for frequency-dependent demodulation of photothermal signals. The in-phase (X) and quadrature (Y) components are then output and recorded by a DAQ system. Through vector subtraction of AC components, OPTM images are generated to provide photothermal phase gradient (PTPG) information. The absorption images are generated by a vector summation operation for comparison with PTPG images.

### OPTM imaging of microparticles in a scattering medium

To validate the principle of oblique photothermal detection, we employed the OPTM to image well-defined microparticles in a scattering medium. Specifically, 10-µm PMMA beads were embedded in a scattering medium composed of 1% intralipid and PDMS mixture, then imaged using the OPTM (**Figure 3**).

**Figure 3.**
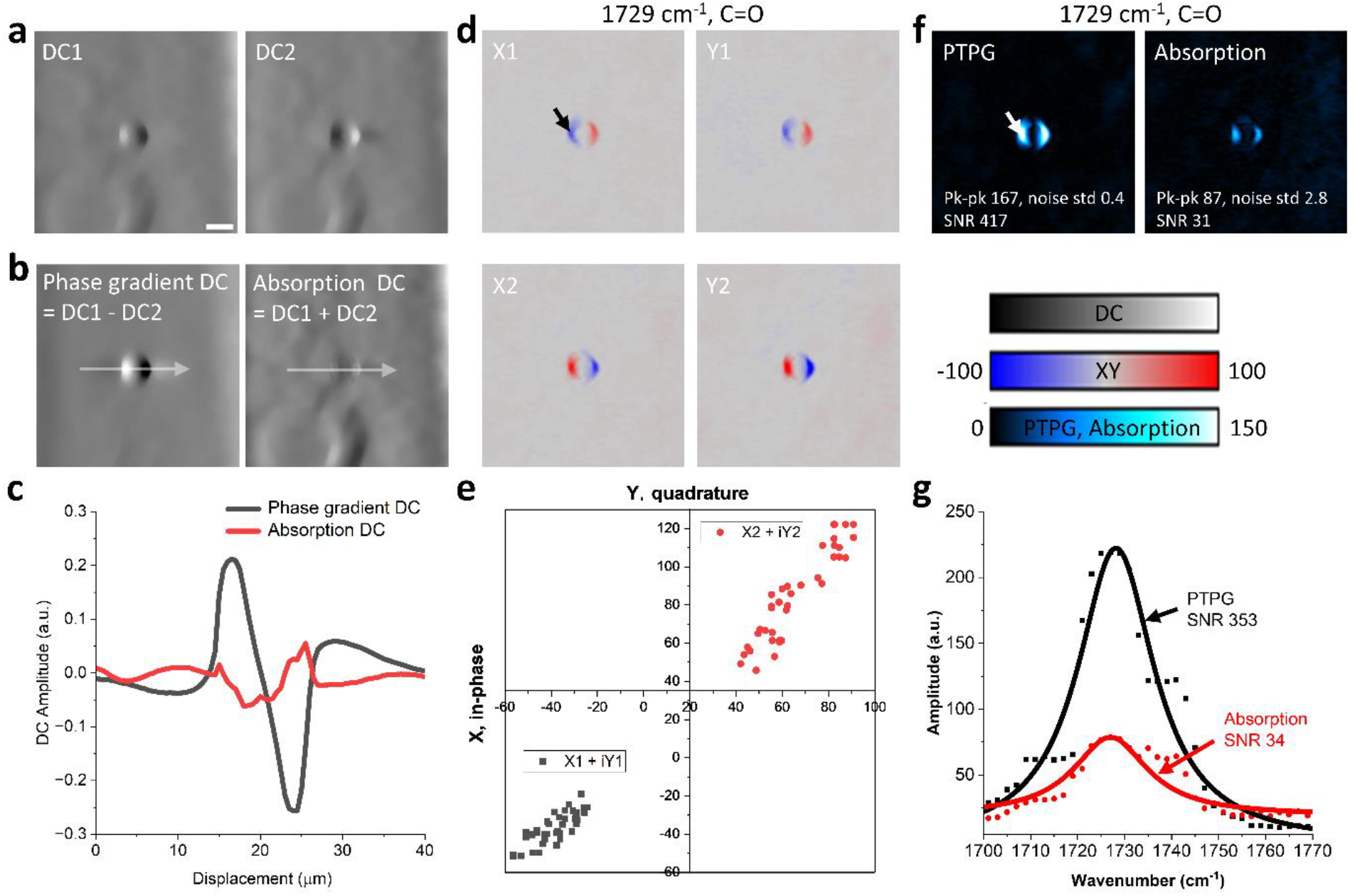
OPTM imaging of microparticles in a scattering medium. 10-µm PMMA beads was embedded in a scattering medium and imaged by OPTM. **a**, DC images of PD1 and PD2. **b**, phase-gradient and absorption DC images. **c**, line plotted indicated by the white arrows in (**b**). **d**, in-phase (X) and quadrature (Y) images of each photodiode from the lock-in amplifier. **e**, scatter plotted in region of interest indicated by black arrow in (**d**). **f**, photothermal phase gradient (PTPG) and absorption images. **g**, spectra at position indicated by the white arrow in (**f**). Scale bar 10 µm. Field of view 75 × 75 µm^2^. Probe power on sample was 10 mW. IR power on sample was ∼2 mW with repetition rate 390 kHz, pulse width 80 ns. The imaging speed was 2.5 frame per second with a pixel dwell time of 10 µs. SNR: signal-to-noise ratio. Std: standard deviation.

In the DC images, both halves of the split detector yield the phase gradient information of the microparticles (**Figure 3a**). The DC signals show an inversed polarity at the two ends of particles. The DC1 and DC2 images show the inversed phase gradients, akin to taking the first derivative of the refractive index from opposite horizontal directions. By subtracting and summing the DC1 and DC2 images, phase gradient DC and absorption DC images are generated (**Figure 3b**). **Figure 3c** shows the plots of white lines in Figure 3b to illustrate that the phase gradient measurement improves the contrast for visualizing particles in a scattering medium compared to absorption measurements. Because PMMA microparticles have negligible absorption at the probe wavelength of 532 nm, no visible features appear in the absorption DC images.

The photothermal images are acquired with the IR wavenumber of 1729 cm^−1^, contributed by the stretching absorption of C=O bond in PMMA particles (*17*). In the photothermal AC images, a lock-in amplifier demodulates frequency-dependent photothermal signals and generates both in-phase (X) and quadrature (Y) components of each split detector to form a complex photothermal response (X+iY) (**Figure 3d**). The X and Y channels form a vector signal, where X is the real part and Y is the imaginary part. **Figure 3e** shows the vector signals of the split detector in a 2D coordinates from the region of interest indicated by the black arrow in Figure 3d. The vector signal of PD1 (X1+iY1) has an inversed direction compared to the vector signal of PD2 (X2+iY2). This indicates that the photothermal modulations of both photodiodes have opposite signs. Compared with the steady status of the IR pump off, with IR pump heating, one photodiode receives more photons while the other receives fewer. These results confirm the principle of oblique photothermal detection. Subsequently, through a post-processing step involving vector subtraction and summation, we derive photothermal phase gradient (PTPG) images and absorption images (**Figure 3f**). In our results, the signal-to-noise ratios (SNR) of PTPG and absorption images reach 417 and 31, which indicates that OPTM is capable of imaging PMMA particles in a scattering medium with a 13-fold improvement of SNR over absorption image and 7-fold reduced laser noise. Additionally, we acquired the off-resonance images at 1770 cm^−1^ to demonstrate the bond-selectivity of OPTM imaging (**Figure S2**). **Figure 3g** shows the spectra of PTPG and absorption at the region of interest indicated with the white arrow in **Figure 3f**. It indicates that OPTM enhances the spectroscopic intensity of particles in a scattering medium by over 10-fold in comparison with absorption measurement. Together, our results validate the principle of oblique photothermal detection and prove that phase gradient is a sensitive readout of photothermal contrast.

### OPTM unveils two types of lipids in adipocytes under live mouse skin

Different lipid compositions are important indicators for skin cancer, such as, sebaceous carcinoma (*44*). Coherent Raman scattering microscopy has allowed imaging of total amount of lipids using C-H vibrational signals (*1-3*). Here, we demonstrate that OPTM can differentiate free fatty acid and lipid ester in individual adipocytes under live mouse skin. To compare the sensitivity of OPTM versus MIP microscope, we imaged adipocyte cells in the back skin of a live mouse using OPTM and MIP. Adipocytes on the epidermal layer of animal skin are vital to health as they form a barrier against external pathogens and serve as indicators of skin biology and disease (*45*). High-sensitivity adipocyte imaging in live animals can facilitate the study and diagnosis of health status.

We employed the OPTM and the MIP microscope to image the same region in the epidermis layer of a live mouse’s back skin at a penetration depth of 50 µm. To differentiate different lipid contents, multispectral OPTM and MIP images were acquired from 1700 to 1770 cm^−1^ with a step size of 2 cm^−1^, which is contributed by C=O vibration absorption of lipids (*46-48*) (**Movie S1**). The DC images of OPTM and MIP were acquired simultaneously, measuring phase gradient, absorption, and back scattering information (**Figure S3**).

Following the flowchart in **Figure 2e**, photothermal phase gradient images were acquired by OPTM. Meanwhile, absorption images were generated for a comparison purpose. MIP images were acquired independently using pinhole filtering approach as illustrated in Figure S1. The photothermal images at two IR wavenumbers, 1704 and 1742 cm^−1^, are selected to present the fatty acid and lipid ester-rich distributions (**Figure 4a-c**). The absorption peaks of the fatty acid and lipid ester are contributed by the vibrational absorption of acidic and esterified C=O bond in lipids, which are consistent with FTIR spectroscopy measurement in animal skin samples (*46-48*). Off-resonance images at wavenumber 1770 cm^−1^ were acquired to confirm bond-selective imaging of OPTM and MIP. Spectra in two regions of interest in **Figure 4a** are plotted to illustrate the absorption peaks of fatty acids and lipid esters at IR wavenumbers 1704 and 1742 cm^−1^ (**Figure 4d**).

**Figure 4.**
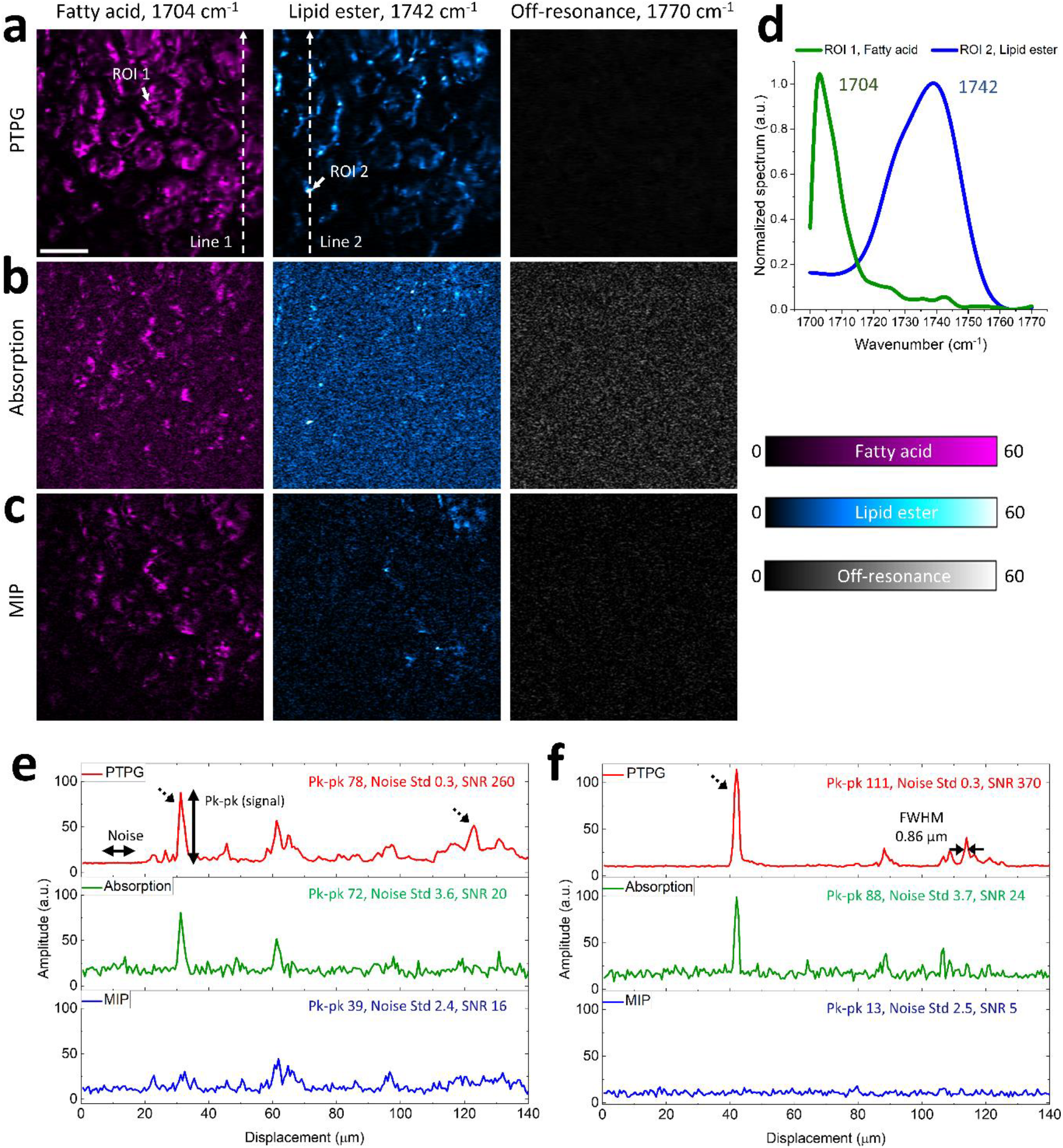
OPTM unveils two types of lipids in adipocytes under live mouse skin. OPTM and MIP imaging of adipocyte cells in the back skin of a live mouse at a penetration depth of 50 µm. Photothermal phase gradient (PTPG), (**b**) absorption, and (**c**) MIP images were acquired in a same field of view. **d**, PTPG spectra of two regions of interest in (a) illustrating the absorption peaks of the fatty acid and the lipid ester. (**e**) Line 1 and (**f**) line 2 plotted in the fatty acid and the lipid ester images. Scale bar 30 µm. Field of view 150×150 µm^2^. Probe power on sample was 10 mW. IR power on sample was ∼2 mW with repetition rate 390 kHz, pulse width 80 ns. The imaging speed was 1.6 frame per second with a pixel dwell time of 10 µs.

To quantify the sensitivity improvement of OPTM over MIP, we plotted intensities along two dash lines in the fatty acid and lipid ester images (**Figure 4e&f**). Regarding the noise levels, PTPG signals show an 12-fold and 8-fold suppression of noise compared to the absorption signals and the MIP signals, respectively. These results indicate that, OPTM can effectively suppress the common-mode noise, primarily contributed by laser intensity noise, via a balanced detection strategy. In parallel to noise suppression, OPTM enhances the photothermal signal by up to 8-fold over MIP owing to the higher photon collection efficiency. As indicated in dashed black rows in **Figure 4e&f**, OPTM enables visualization of lipid structures not detectable in MIP. The peak at the PTPG image of lipid ester has a full width at half maximum of 0.86 µm, indicating sub-micron lateral resolution of OPTM.

### Depth-resolved in vivo OPTM imaging of mouse skin

Depth-resolved infrared spectroscopic imaging is essential for accurately visualizing and analyzing chemical contents in the intricate layers and structures within complex tissues (*49*). Despite its importance, in vivo infrared depth-resolved imaging remains an unmet need in the field. Our OPTM technique addresses this gap by offering optical sectioning capability at visible light resolution through a pump-probe approach.

**Figure 5** demonstrates in vivo depth-resolved OPTM imaging. We utilized OPTM to image the ear of a live mouse at different penetration depths. At each depth, protein images were acquired at an IR wavenumber of 1553 cm^−1^, contributed by the absorption of the Amid II band (*8*). Two lipid contents, fatty acid and lipid ester, were imaged at IR wavenumbers of 1704 and 1742 cm^−1^ due to the vibrational absorption of acidic and esterified C=O bond in lipids (*46-48*). Additionally, off-resonance images at 1770 cm^−1^ were acquired to confirm bond-selectivity of OPTM imaging.

On the stratum corneum (SC) layer at a depth of 10 µm beneath skin surface, which is the outmost layer of skin, the mouse hair is visualized (**Figure 5a**). SC shows sparse lipid and protein distributions because it includes mostly of dead cells. The viable epidermis layer, right beneath the SC layer, mainly consists of live cells providing nutrition to the skin. Thus, the protein and lipid content are uniformly distributed at a depth of 50 µm (**Figure 5b**). As the penetration depth increases to 75 µm into the epidermis layer, sebaceous glands become visible, predominantly composed of lipid esters and some fatty acids (**Figure 5c**). At a penetration depth of 100 µm, which is equal to one mean free length of 532-nm photons (*50*), the phase gradient DC image shows a good contrast for resolving deep tissue features (**Figure S4**). Photothermal phase gradient images still show differentiable chemical distributions. Thus, we determine the penetration depth limit of OPTM to be 100 µm for imaging animal’s skin. Our results collectively validate OPTM’s depth-resolved imaging capability in live animals.

**Figure 5.**
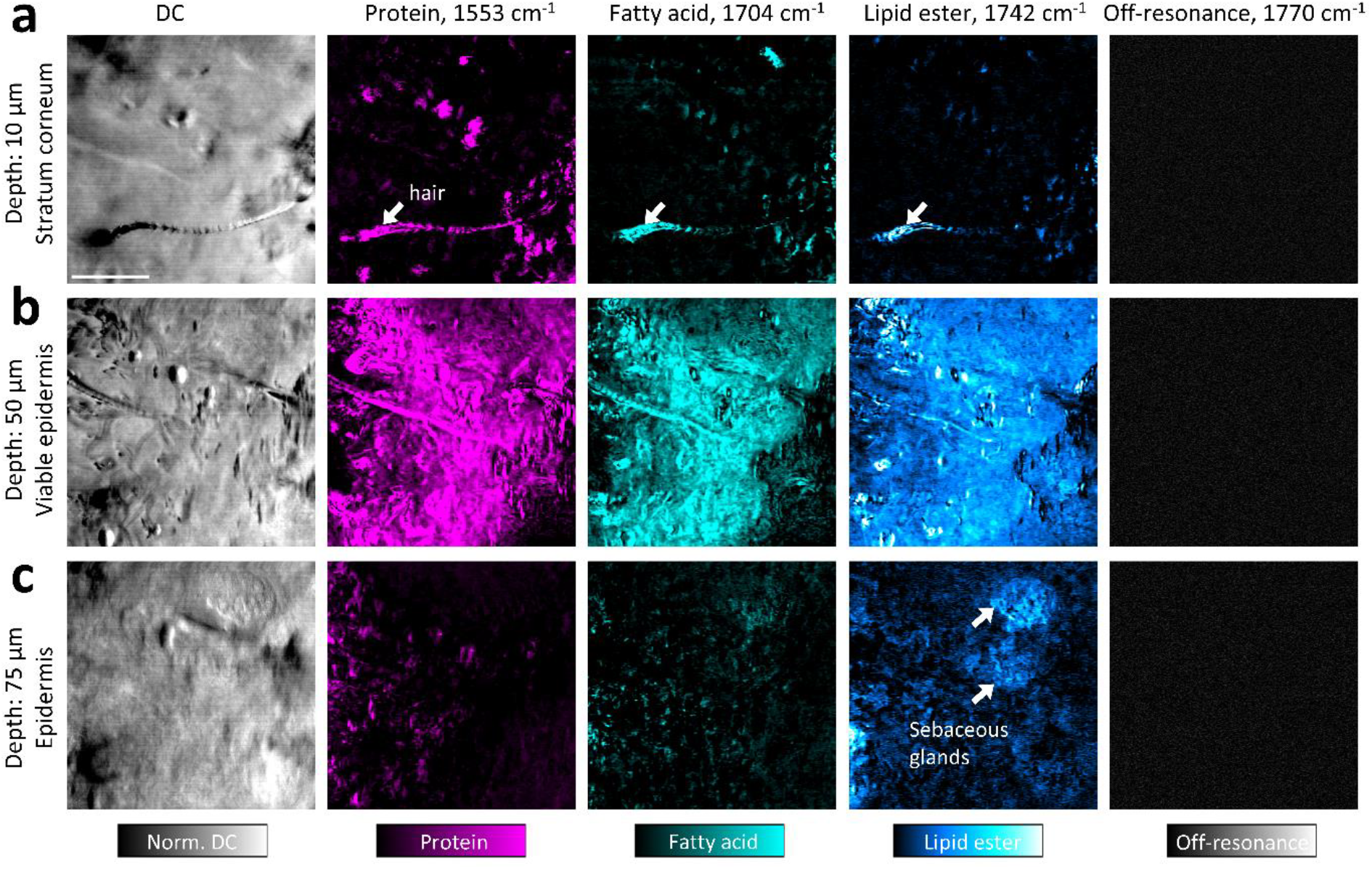
Depth-resolved in vivo OPTM imaging of mouse skin. Phase-gradient DC images and photothermal phase gradient images at imaging depths of (**a**) 10 µm (stratum corneum layer), 50 µm (viable epidermis layer), and (**c**) 75 µm (epidermis layer). Scale bar 50 µm. Field of view 160×160 µm^2^.

### In vivo investigation of topical drug pathway inside mouse skin

Topical transdermal drug delivery presents a promising alternative to both oral administration and hypodermic injections (*51*). In vivo investigation of topical drug delivery inside animal skin facilitates analysis of the drug pathway, guiding the development of new biomedicine (*52*). Quantitative evaluation of effective transdermal drug delivery is crucial to optimize therapeutic efficacy while minimizing systemic side effects (*53*). In **Figure 6**, we demonstrate that OPTM facilitates high-resolution and depth-resolved in vivo tracking of topical drug pathways in mouse skin with quantitative evaluation capabilities. Specifically, we explore the skin penetration behavior of benzoyl peroxide (BPO), a commonly used active pharmaceutical ingredient. BPO is particularly effective for acne treatment, as it penetrates hair follicles to exert antimicrobial and keratolytic effects (*54*).

**Figure 6.**
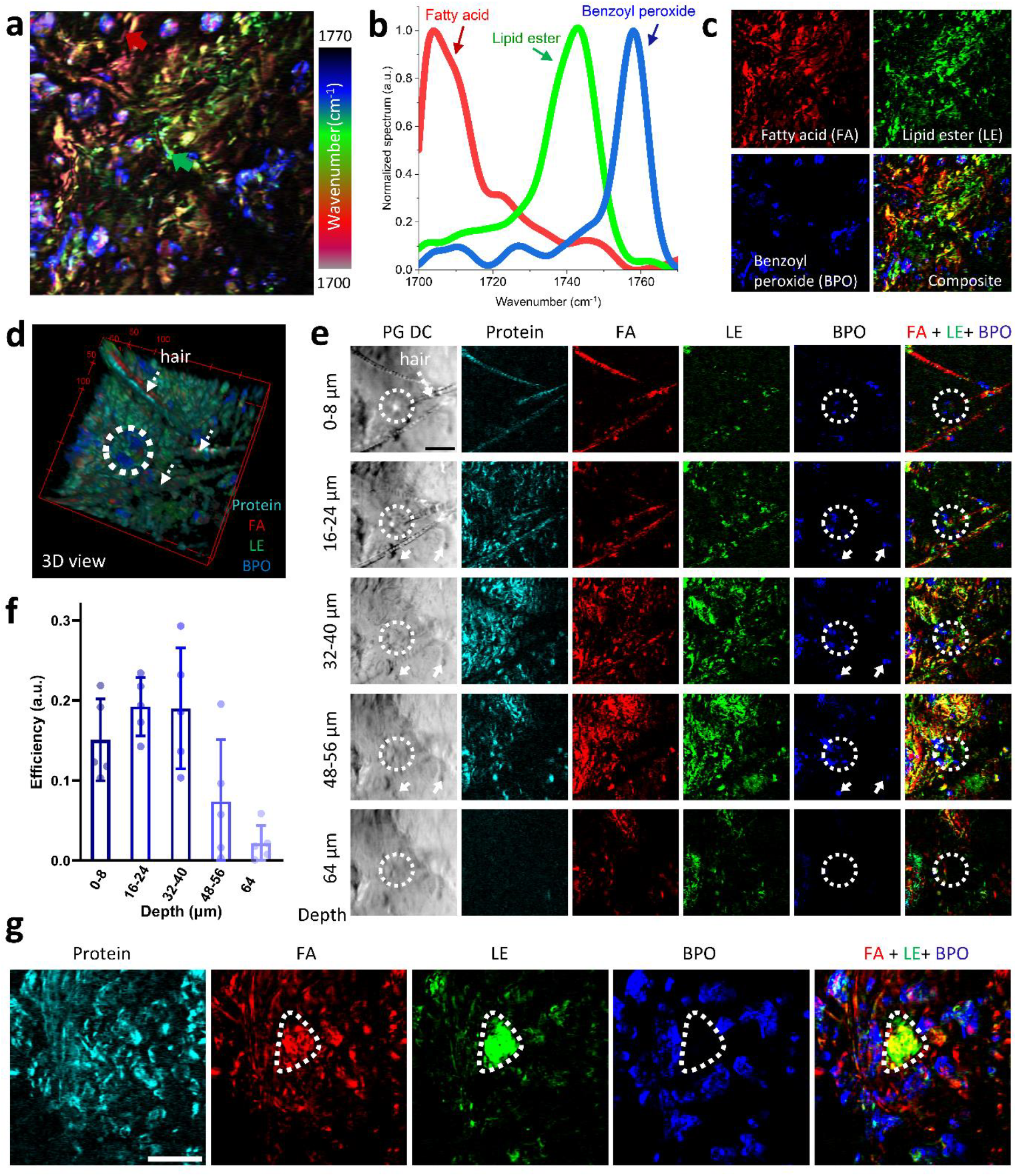
In vivo investigation of topical drug pathway inside mouse skin. **a**, Pseudo-color hyperspectral OPTM images of skin of a live mouse at a depth of 32 µm with benzoyl peroxide administrated. **b**, reference spectra of fatty acid, lipid ester, and benzoyl peroxide. **c**, images of fatty acid, lipid ester, and benzoyl peroxide after applying LASSO spectral unmixing method to the hyperspectral image (a) to differentiate different contents. **d**, a captured 3D view of Movie S2 to illustrate mouse skin chemical structure and topical BPO distribution. **e**, phase gradient DC images and PTPG images at different penetration depths in mouse skin. **f**, efficiency of topical BPO delivery at different depths, based on analysis of 5 FOV images shown in Figure 6e and S7. **g**, endogenous chemical structure and topical BPO distribution within the skin layers at a depth of 32-48 µm, where the sebaceous glands are located. Additional images of the same region at different depths are presented in Figure S8. Scale bar 50 µm. Field of view 180×180 µm^2^. Probe power on sample was 10 mW. IR power on sample was ∼2 mW with repetition rate 390 kHz, pulse width 100 ns. The imaging speed was 1.1 frame per second with a pixel dwell time of 10 µs. FOV, field of view. FA, fatty acid. LE, lipid ester. BPO, benzoyl peroxide.

To visualize both BPO and endogenous chemical contrasts, we performed multispectral in vivo OPTM imaging at various depths in mouse skin, following the administration of 5-µL BPO samples. **Figure 6a** shows a pseudo-colored multispectral image acquired at a depth of 32 µm beneath the mouse skin surface. To quantitatively disentangle the different chemical components from the composite spectrum, we employed a least absolute shrinkage and selection operator (LASSO) for spectral unmixing (*55*). In LASSO, it is essential to have reference spectra of different chemical contents. As illustrated in **Figure 6b**, the reference spectra of fatty acid and lipid ester are acquired from the regions indicated by the red and green arrows in **Figure 6a**, which are rich in fatty acid and lipid ester, respectively. The reference spectrum of BPO is acquired from the infrared photothermal spectrum of drug samples (**Figure S5**). **Figure 6c** shows the spectral unmixing results of LASSO that separate the fatty acid, lipid ester, and administrated BPO contents from the raw multispectral image. An overlap of three chemical components shows the relative distributions of endogenous lipid and the drug contents. LASSO spectral unmixing was applied to other depth-resolved imaging data to trace topical BPO pathway. A control experiment, conducted without drug administration, shows no visible contrast in the BPO channels (**Figure S6**). Through depth-resolved infrared spectroscopic imaging and LASSO spectral unmixing, we demonstrate OPTM enables three-dimensional visualization of chemical content in mouse skin, illustrating endogenous skin chemical structure and administrated BPO distribution (**Movie S2 & S3**).

**Figure 6d** shows a captured 3D view in Movie S2, in which the mouse hairs and the hair follicles opening can be seen. Protein images were acquired at 1553 cm^−1^ contributed by the absorption of Amid-II band. To unveil the BPO pathway under mouse skin, we show the depth-resolved phase gradient DC and different chemical images illustrating the skin morphology, chemical structure, and drug/skin distributions at different skin layers (**Figure 6e**). At the stratum corneum layer with a depth 0-8 µm, which is the skin surface, the hair follicle locates at one end of the mouse hair as indicated by white dash circle in the phase gradient DC image. There is topical BPO content overlapping with the hair follicle opening. At the epidermis layer with depths of 16-24, 32-40, and 48-56 µm, BPO content overlaps with the hair follicles indicated by white dash circles. It confirms that BPO penetrate into the hair follicles in the skin through its openings, as the acnes happen in the hair follicles (*56*). At a deeper penetration depth of 64 µm, the endogenous chemical and BPO images still have visible contrast, demonstrating the capability of OPTM to trace drug molecules in deep skin layer. Through all skin layers, the distribution of administrated drug content is not uniform but aggregates. Surprisingly, we found that BPO content also penetrates sweat pores as indicated by the solid white arrows in **Figure 6e**. To quantitatively evaluate the delivery efficiency of topical BPO, we took another four group images at different field of views (**Figure S7**). Since BPO is effective to treat acne by penetrating hair follicles to perform antimicrobial and keratolytic activities, we quantify the delivery efficiency by computing the ratio between the amount of drug content within hair follicles and within the whole field of view at a same depth (**Figure 6f**). From the skin surface, with a deeper penetration depth, the BPO content within the hair follicle increases at first, then followed by a decreasing. It indicates that topical BPO enters the hair follicles through its opening and remains inside the follicles. The delivery efficiency of BPO is less than 30% in skin layers. Such a low efficiency indicates the treatment of skin acne needs a long-term administration of topical BPO, which is aligned with previous studies of acne treatment by BPO (*57*).

The interaction between topical BPO and sebaceous glands has not been clarified, and the side effects of BPO remain controversial (*58-60*). Here, OPTM is used for monitoring the spatial distributions of BPO and sebaceous glands. **Figure 6g** shows the skin layers, including the sebaceous glands, at a depth of 32-48 µm within the volume shown in **Movie S3**. The sebaceous glands are visualized at endogenous lipid images indicated by white dash circles, which does not overlap with BPO. **Figure S8** shows the depth-resolved images in the same region for illustrating the drug pathway within the skin layers.

In summary, OPTM shows that BPO penetrates the mouse skin through hair follicles opening, sweat pores, and other skin areas. The low efficiency of BPO delivery via hair follicles, which is less than 30%, indicates a need for long-term administration of BPO for the treatment of skin acne. In addition, our observations unveil that topically applied BPO does not reach sebaceous glands. Collectively, OPTM enables in vivo depth-resolved tracking of topical drug pathway with quantitative evaluation of delivery efficiency, providing guides to the development of new transdermal biomedicine.

### In vivo OPTM imaging of human skin

Non-invasive chemical analysis of live human subjects plays a crucial role in biomedical research and clinical diagnostics (*61*). Providing fingerprint information with high spatial resolution, OPTM is a good candidate for non-invasive chemical analysis of human skin. In OPTM, the use of visible probe offers sub-micron resolution, while the improved sensitivity allows low-dose IR spectroscopic imaging without the concern of photodamage. **Figure 7** demonstrates OPTM’s ability to non-invasively image the metabolic markers under live human skin and the topical transdermal drug content on the skin surface.

**Figure 7.**
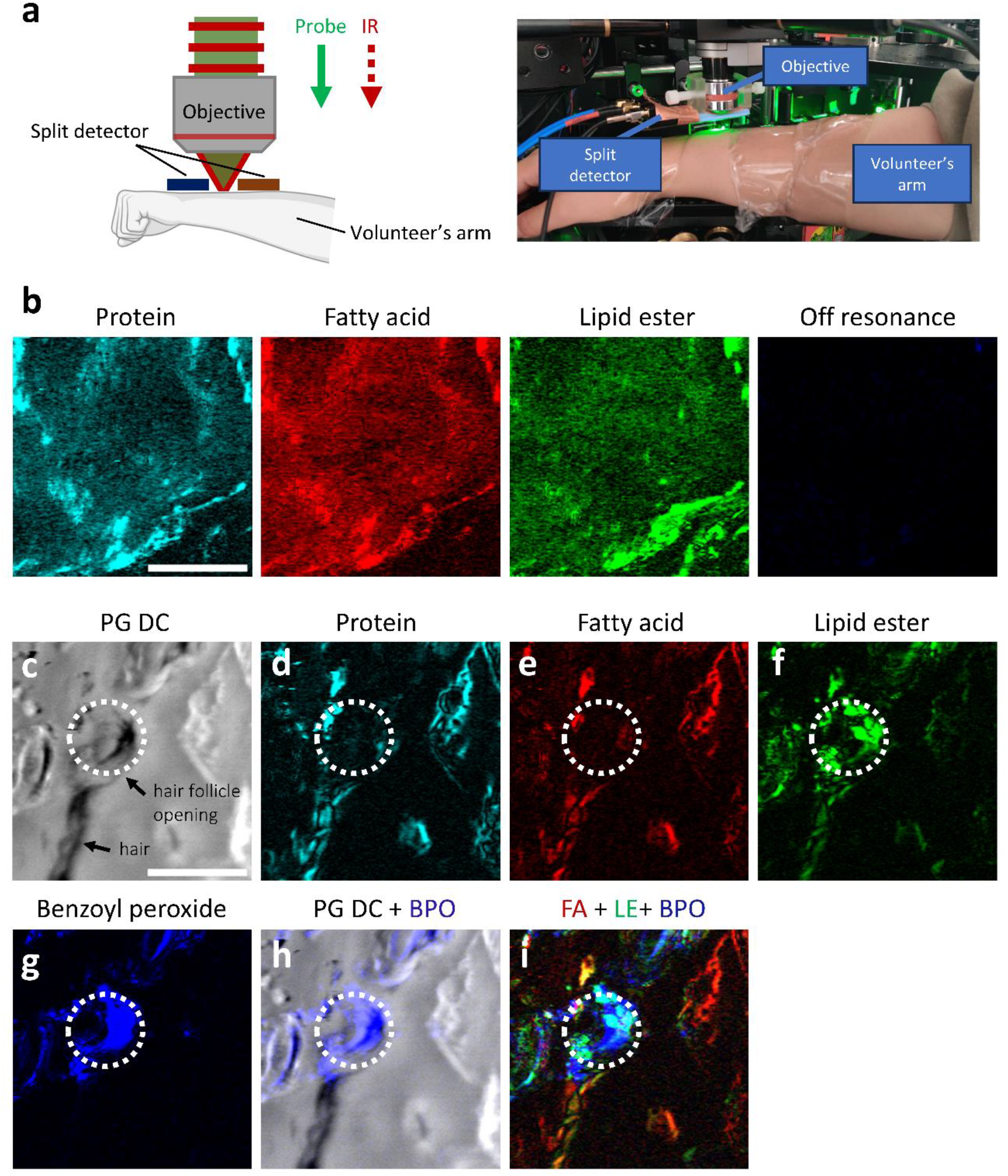
In vivo OPTM imaging of human skin. **a**, schematic and photo of in vivo OPTM imaging of a human’s forearm skin. **b**, photothermal phase gradient images of endogenous proteins and lipids in the viable epidermis layer with a depth of 40 µm without topical drug administration. **c**, phase gradient DC image of the stratum corneum showing a hair on the skin surface and the hair follicle opening. **d-g**, photothermal phase gradient images of protein (**d**, 1553 cm^−1^), fatty acid (**e**, 1704 cm^−1^), lipid ester (**f**, 1742 cm^−1^), and benzoyl peroxide (**g**, 1760 cm^−1^). **h**, an overlap image of phase gradient DC and drug distribution. **i**, an overlap image of endogenous lipid and administrated drug contents. FA, fatty acid. LE, lipid ester. BPO, benzoyl peroxide.

A volunteer’s forearm was imaged by OPTM to visualize the skin chemical structure and the topical drug content upon the skin surface (**Figure 7a**). In **Figure 7b**, OPTM visualizes endogenous proteins and lipids in the viable epidermis layer with a depth of 40 µm without topical drug administration. Off resonance imaging at 1780 cm^−1^ confirms the bond-selective imaging of human skin. The relative distribution of protein and different lipid contents under skin surface are an important health indicator, such as skin aging (*62*) and inflammatory skin disease (*63*).

Additionally, we demonstrate that OPTM enables to trace the topical drug pathway on human skin surface. **Figure 7c-g** show phase gradient DC and PTPG images of endogenous chemicals and administrated benzoyl peroxide contents on human arm skin with a depth of 8 µm. From phase gradient DC image in **Figure 7c**, the hair follicle opening located at the end of one human hair can be seen. **Figure 7g** shows that topically administrated BPO contents on human skin surface is also not uniform but aggregates, which aligns with the observations of mouse imaging in **Figure 6**. The BPO content overlap with hair follicles opening is shown in **Figure 7h**, which demonstrates that BPO penetrates skin through hair follicle opening and other skin areas. The percentage of BPO through follicle is estimated to be 46.5% based on two different field of views.

**Figure 7d-f** show a mixture of chemical content within the hair follicles, such as sebum. **Figure 7i** shows the overlap of endogenous chemicals and topical BPO, demonstrating that OPTM allows to map both topical biomedicine and skin chemical structure. To verify the imaging fidelity, a control experiment was performed by using OPTM to image human arm skin without BPO administration (**Figure S9**). The BPO image in the control group shows no visible contrast within the hair follicles. It confirms that the contrast of **Figure 7g** comes from the administrated BPO. Collectively, we demonstrate that OPTM allows non-invasive infrared spectroscopic imaging of human skin in vivo. OPTM visualizes the metabolic biomarker beneath human skin surface, facilitating heath diagnosis in clinical application. OPTM also allows monitoring of topical drug content on human skin, facilitating the development of transdermal biomedicine.

## Discussion

Oblique photothermal detection challenges the conventional wisdom of photothermal sensing by fundamentally altering the photon collection strategy. Traditional photothermal methods typically measure a small fraction of backward-scattered photons filtered through an iris, which inherently limits the number of collected photons. This limitation arises because only photons scattered in specific directions are detected, reducing the signal intensity and detection sensitivity. In contrast, oblique photothermal detection employs a split detector positioned on the sample surface, enabling the collection of both backward and forward-scattered photons. The forward-scattered photons are redirected into the backward direction through multiple scattering events within the sample. Essentially, this method significantly enhances the photon collection efficiency.

Furthermore, OPTM suppresses the laser noise via balanced detection. By leveraging the inversed photothermal modulations on the split detector, OPTM amplifies the photothermal signal amplitude by analyzing the differential response between the two halves of the split detector. Since the photothermal signal predominantly occupies the low-frequency region due to its low-duty cycle nature, traditional photothermal detection methods is compromised by 1/f laser noise. OPTM collects 500 times more photons from tissue, and mitigates the laser noise through a subtraction operation akin to the balanced detection strategy (*39*).

OPTM could detect weak scatterers in scattering samples by measuring the photothermal phase gradient of the object rather than back scattering signals. The amount of back scattering photons from the object is limited by the minimal refractive index difference within tissue (*64*). Traditional photothermal detections, which also gather other diffuse and specular reflected photons, could have the scattering signals from the focus overwhelmed by the laser noise. Oblique photothermal detection addresses this challenge by utilizing a split detector near sample surface to acquire phase gradient as an ultrasensitive readout of photothermal signals. By measuring the photothermal phase gradient rather than back scattered photons, OPTM boosts the imaging sensitivity, enabling visualization of weak scatterers in complex tissue. Though we have focused on skin applications, OPTM is applicable to other tissues. **Figure S10** demonstrates that OPTM facilitates highly sensitive IR spectroscopic imaging of a mouse brain slice. OPTM allows for in situ molecular analysis within the soma and blood vessels, revealing the potential for detailed molecular profiling in complex tissues.

There is potential for further enhancing OPTM’s penetration depth. The absorption of IR photons and the scattering of probe photons are two main factors of OPTM to penetrate deeper. However, the attenuation length of IR photons (< 200 µm) is still larger than the attenuation length of 532-nm probe photons (100 µm) (*13, 50*). Thus, the main limiting factor is the strong scattering of probe photons within tissue due to the short visible wavelength used. Employing a longer probe wavelength could facilitate deeper imaging by reducing optical scattering (*50*).

In summary, by providing reduced laser noise, enhanced imaging contrast, and improved sensitivity, OPTM enables high-sensitivity infrared spectroscopic imaging of chemicals in live animals and human subjects with sub-micron resolution. OPTM opens translational opportunities for molecule-based clinical diagnosis and study of topical drug delivery through human skin. As a platform technology, OPTM is not limited to the mid-infrared window and can be broadly adapted to other photothermal microscopy techniques such as overtone photothermal microscopy (*65-67*) and visible photothermal microscopy (*68*).

## Materials And Methods

### Monte Carlo simulation

A Monte Carlo simulation was conducted to explore the propagation path of 4-million photons in a uniform scattering layer (*41, 42*). The sizes of the layer were (width) 25 mm, (length) 25 mm, and (depth) 10 mm. A focused beam with a numerical aperture (NA) of 0.5 was launched on the surface of simulation layer. The NA of simulated focused beam is equal to the NA of reflective objective used in the OPTM and MIP microscope. The scattering coefficient µ_s_ was set from 20 to 125 cm^−1^, referring to the attenuation length (*L* = 1⁄μ_*s*_) of 500 to 80 µm. The absorption coefficient was set to 0.1 cm^−1^.

### Oblique photothermal microscope

Our OPTM (**Figure 2**) couples a 532-nm continuous wave probe laser (Samba, HUBNER photonics) with a pulsed quantum cascade laser (MIRcat 2400, Daylight Solutions) tunable from 900 cm^−1^ to 2,300 cm^−1^ via a Germanium window (GEBBAR-3-5-25, Andover Corporation). The visible/IR combined beam is directed to a 2D galvo scanner (Saturn 5B, ScannerMax) and then through a reflective relay optics, primarily comprising a concave mirror with a 150-mm focusing length (CM1, CM254-150-P01, Thorlabs) and a concave mirror with a 250-mm focusing length (CM2, CM254-250-P01, Thorlabs). Subsequently, both beams are focused onto the sample by a reflective objective (LMM40X-P01, Thorlabs). In the detection part, a split detector consisting of two identical photodiodes (S3994-01, Hamamatsu or FDS1010, Thorlabs) is biased with a 100-volt voltage to elevate its saturation threshold. A 402 kΩ resistor (R) and a 0.1 µF capacitor (C) form an integrated RC low-pass filter to reject high-frequency noise from the biased voltage supply. A lab-designed printed circuit board integrates the photodiodes and other electrical components (**Figure S11a**). The split detector is mounted on a homebuilt printed circuit board. A copper mesh (PSY406, Thorlabs) is used as a cover for electromagnetic shielding (**Figure S11b**). Here, a 2-mm slit width between two photodiodes is created to allows the combined pump and probe beam to pass through. A 3D printed holder is glued at the other side of PCB board so that it can be mounted on the imaging objective (**Figure S11c**). The output of each photodiode is then split into DC and AC signals using a biased tee (ZFBT-4R2GW+, Mini-circuits). The DC signals are collected by a low-noise data acquisition card (DAQ) (Oscar, Vitrek), while the AC signals are amplified by a RF low-noise amplifier (CMP61665-1, Gain 40 dB, NF Corporation). Then, the amplified AC signals are fed into a lock-in amplifier (HF2LI, Zurich) for frequency-dependent demodulation of photothermal signals. A multichannel DAQ board (National Instruments, PCIe-6363) is used for real-time data acquisition.

### PMMA microparticles in a scattering medium

Polydimethylsiloxane (PDMS) (Sylgard 184, Dow Corning Corporation) was prepared with a base-to-curing agent ratio of 10:1. Then, 10-µm Polymethylmethacrylate (PMMA) beads (17136-5, Polysciences) were added to the PDMS mixture along with 1% intralipid (I141, Sigma-Aldrich). The mixture was degassed in a vacuum chamber for 30 minutes to remove residual air bubbles. Following this, the composite was then solidified by heating it in an oven at 70°C for 30 minutes.

### In vivo imaging of mouse skin

Male adult C57BL/6J and BALB/c mice were used in this study. The details of mouse preparations for imaging are illustrated in **Figure S12**. In step 1, the mouse was anesthetized with isoflurane, and the hair was removed or shaved to clean the skin surface. Subsequently, the mouse was given a subcutaneous injection of 50-100 mg/kg Ketamine and 5-10 mg/kg Xylazine, enabling the transition to the imaging process. The level of anesthesia was assessed by the loss of response to pinching the base of the tail. In step 2, a biocompatible glue was applied to fix the mouse’s skin to a Teflon block, which served as a scattering substrate and mitigated breathing artifacts during imaging. A warming bag was placed under the mouse to maintain its body temperature. In step 3, the mouse was transferred to the OPTM for imaging. In step 4, after imaging, the mouse skin was detached from the Teflon block without visible hurt. Then, the animals were euthanized via isoflurane anesthesia followed by cervical dislocation. The preparation process for imaging ear skin of mouse is similar to the process described above, excluding the removal of hair.

### In vivo imaging of human skin

To mitigate the motion artefact during imaging, the volunteer’s arm was fixed on a holder by transparent tapes. The protein, fatty acid, lipid ester, and benzoyl peroxide contents were imaged at the IR wavenumber of 1553 cm^−1^, 1704 cm^−1^, 1742 cm^−1^, and1760 cm^−1^, respectively, for shortening imaging time without losing information. No visible damage can be seen on human skin after the imaging experiment.

### Infrared photothermal spectrum of benzoyl peroxide

A skin-care drug was acquired from a CVS drug store which is used to treat skin acne (**Figure S5a**). It includes 10% benzoyl peroxide (BPO) as the active pharmaceutical ingredient (**Figure S5b**). One BPO molecule includes two phenol rings and two C=O bonds (**Figure S5c**). To explore the spectral features of the drug molecule, we used a co-propagation MIP spectroscope to measure the infrared photothermal spectrum of drug (**Figure S5d**) (*17*). The drug sample was sandwiched by a coverslip and a CaF_2_ glass substate which is IR transparent. A photodiode was placed in the forward direction with an iris to measure photothermal signals because most probe photons went through such transparent samples. **Figure S5e** shows the normalized infrared photothermal spectrum of benzoyl peroxide indicating the peak locates at 1760 cm^−1^, which is contributed by vibrational absorption of C=O bond in the BPO molecule (*69*).

### Pixel-wise LASSO algorithm for chemical map unmixing

Least absolute shrinkage and selection operator (LASSO) used in this study unmixes hyperspectral OPTM imaging data into different chemical components (*55*).

In LASSO, given the hyperspectral OPTM dataset registered as spatial domain, *N*_*x*_, *N*_*y*_, spectral domain, *N*_*λ*_, to unmix into multiple chemical components, we first reshape the 3D hyperspectral stack to a 2D matrix 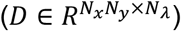 by arranging the pixels in the raster order. Assuming the number of interested chemical channels as *K*, a model is used to decompose the data matrix into the multiplication of concentration maps 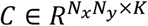 and spectral profiles of pure chemicals 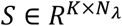

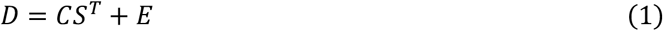

where E is the error. Here, we add a L1-norm regularization to each row of the concentration matrix and solve the original inverse problem in a row-by-row manner through LASSO regression:

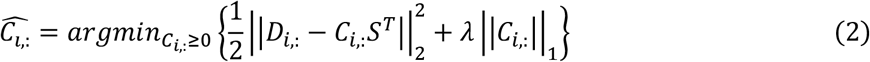

Where *C*_*i*,:_ is a k-element nonnegative vector representing the *i*_*th*_ row of the concentration matrix, 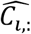 is the output of LASSO regression, *D*_*i*,:_ is the *i*_*th*_ row of the data matrix, *λ* is the hyperparameter that tunes the level of sparsity.

## Acknowledgements

This work was supported by NIH R35 GM136223, R01 EB032391, R01 EB035429, R33 CA261726 to J.-X.C., and in part by grant number 2023-321163 from the Chan Zuckerberg Initiative DAF, an advised fund of Silicon Valley Community Foundation. The authors thank Ling Fang and Jianpeng Ao for help in human imaging; Dashan Dong and Qing Xia for help in refining this paper; Hongjian He for useful discussions; Haonan Lin for providing the LASSO algorithm; and Guo Chen for help in fabrication of detector holder.

## Author contributions

J.-X.C and M.L. conceived the study. J.-X.C and M.L. designed the project and drafted the paper. M.L. performed the experiments and data analysis. X.S. helped in data analysis and made intellectual contributions to experimental design. H.N. helped in circuits design and fabrication. G.D. helped in spectral unmixing. Y.Y. helped in data analysis. C.M. helped in animal operations. J.-X.C and J.M. provided overall guidance to this work.

