## Supplementary Figures for "Ultrasensitive in vivo infrared spectroscopic imaging via oblique photothermal microscopy"

Mingsheng Li et al.

**Supplementary Figures** (Figure S1 to S12)

**Supplementary Videos** (Movie S1 to S3)

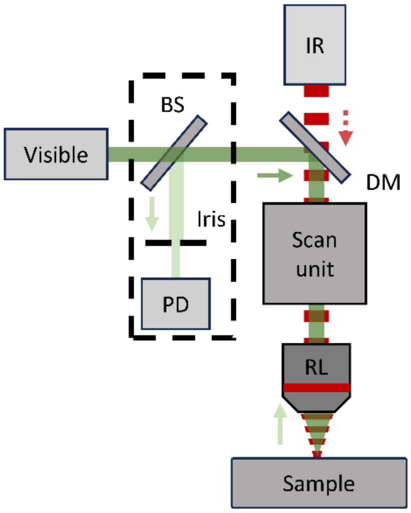


**Supplementary Figure 1**. **Schematic of a mid-infrared photothermal microscope.** In the detection path, a non-polarizing beamsplitter (R50:T50) is used to reflect the backward propagated probe photons, which is filtered by an iris before reaching to a remote photodiode. The signal from photodiode is separated into DC and AC components by a biased tee. The DC component is recorded by a DAQ card. The AC component is amplified by a RF amplifier and delivered into the lock-in amplifier to generate photothermal signals. BS: beam splitter. PD: photodiode. DM: dichroic mirror. RL: reflective objective.

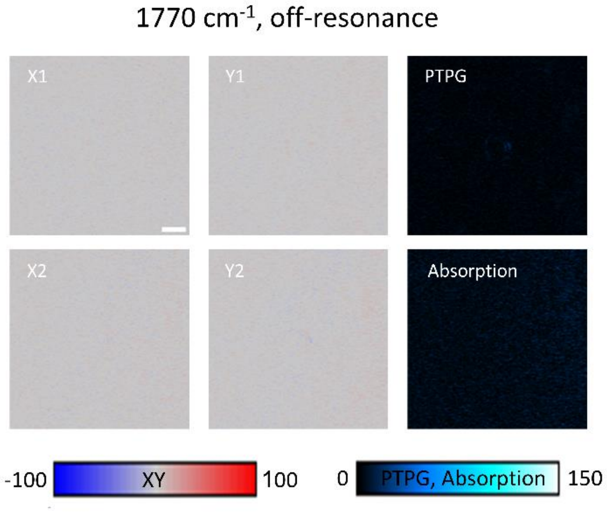


**Supplementary Figure 2. Images acquired by OPTM at off-resonance of infrared absorption (1770 cm$^{-1}$)**. Scale bar 10 μm. Field-of-view 75× 75 μm$^2$. Probe power on sample was 10 mW. IR power on sample was ~2 mW with repetition rate 390 kHz, pulse width 80 ns. The imaging speed was 2.5 frame per second with a pixel dwell time of 10 μs.

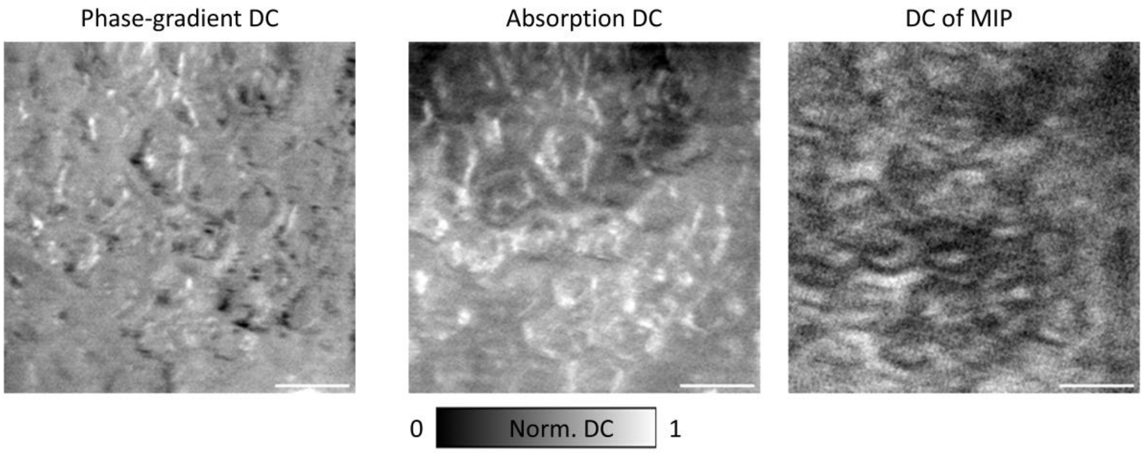


**Supplementary Figure 3.** DC images of phase gradient, absorption, and MIP modalities in Figure 3.

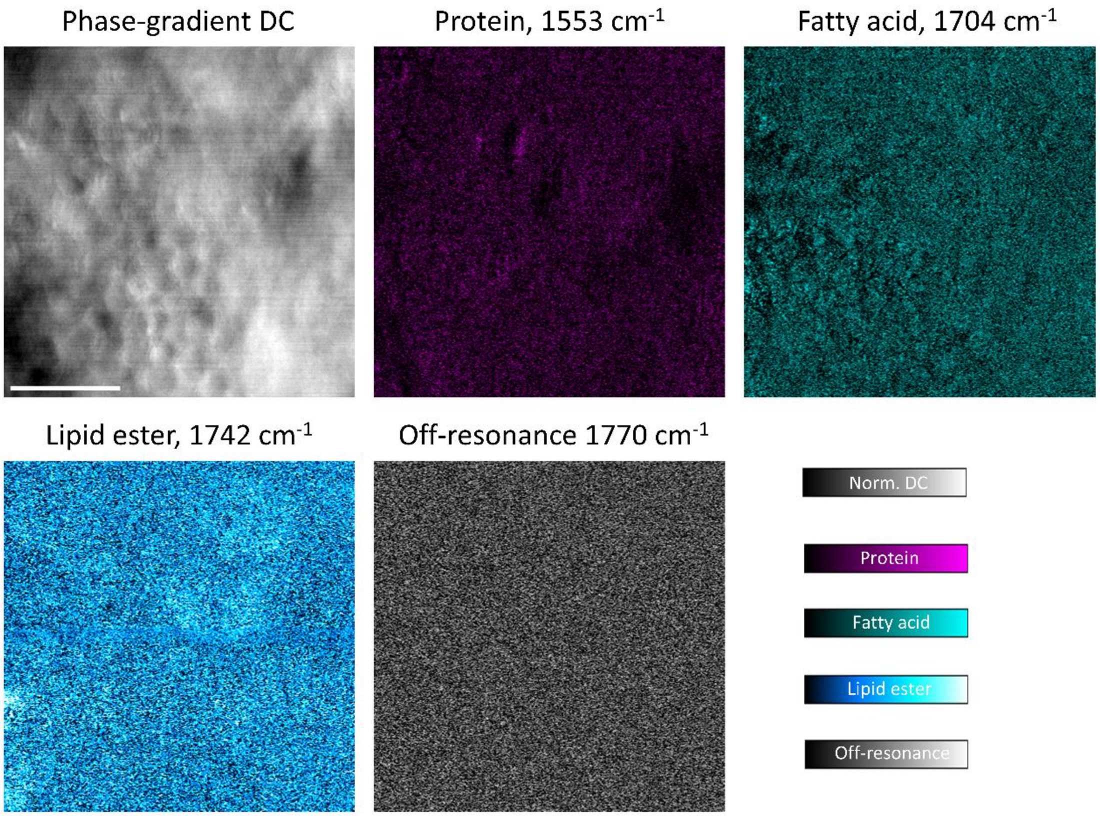


**Supplementary Figure 4. In-vivo OPTM imaging of mouse ear skin at a penetration depth of 100 µm.** Scale bar 50 µm. Field-of-view 160×160 µm$^2$.

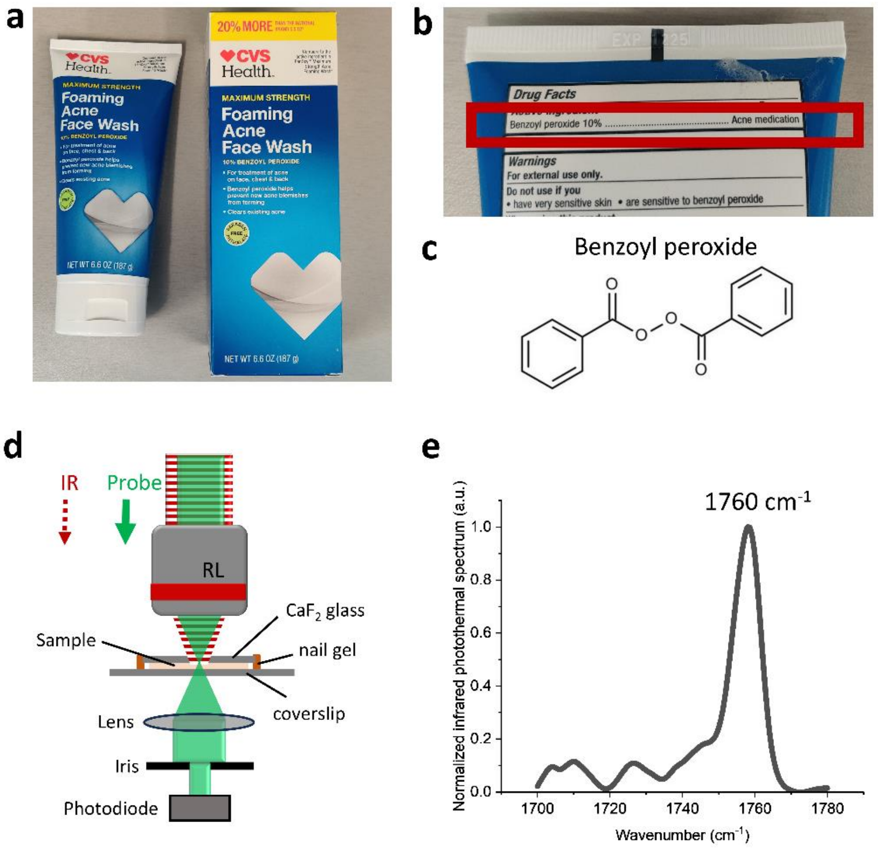


**Supplementary Figure 5. Mid-infrared photothermal spectrum of benzoyl peroxide. a,** photo of skin-care product purchased from CVS. **b,** a close-up photo to indicate that the active ingredient is 10% benzoyl peroxide. **c,** molecular structure of drug molecule, benzoyl peroxide. **d**, experimental set-up for measuring infrared photothermal spectrum of samples. IR, infrared. RL, reflective objective. **e**, normalized infrared photothermal spectrum of benzoyl peroxide gel. The peak at 1760 $cm^{-1}$ is from the esterified C=O bond.

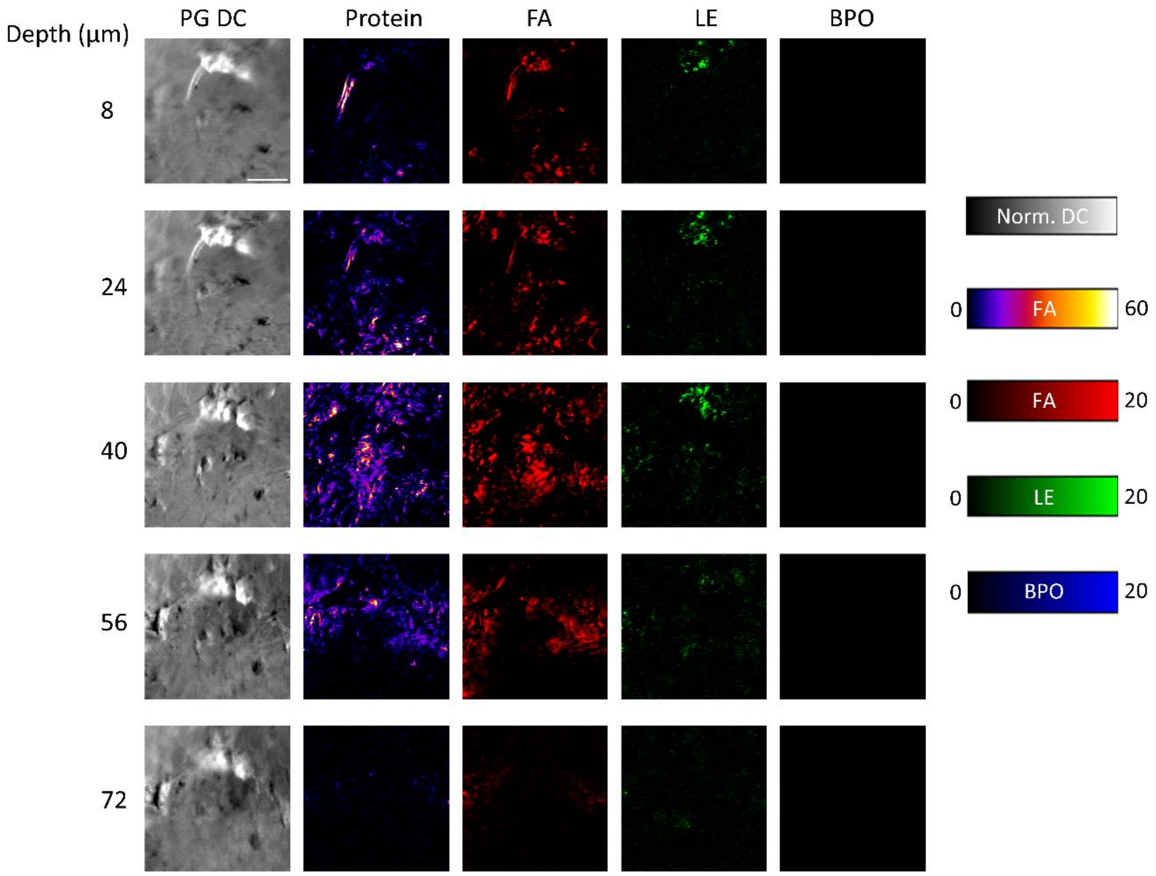


**Supplementary Figure 6. Control group images without benzoyl peroxide administrated**. Phase gradient DC images and PTPG images at different penetration depth in mouse skin. Scale bar 50 μm. Field-of-view 180×180 μm$^2$.Step size 0.6 μm. Probe power on sample was 10 mW. IR power on sample was ~2 mW with repetition rate 390 kHz, pulse width 100 ns. The imaging speed was 1.1 frame per second with a pixel dwell time of 10 μs. FA, fatty acid. LE, lipid ester. BPO, benzoyl peroxide.

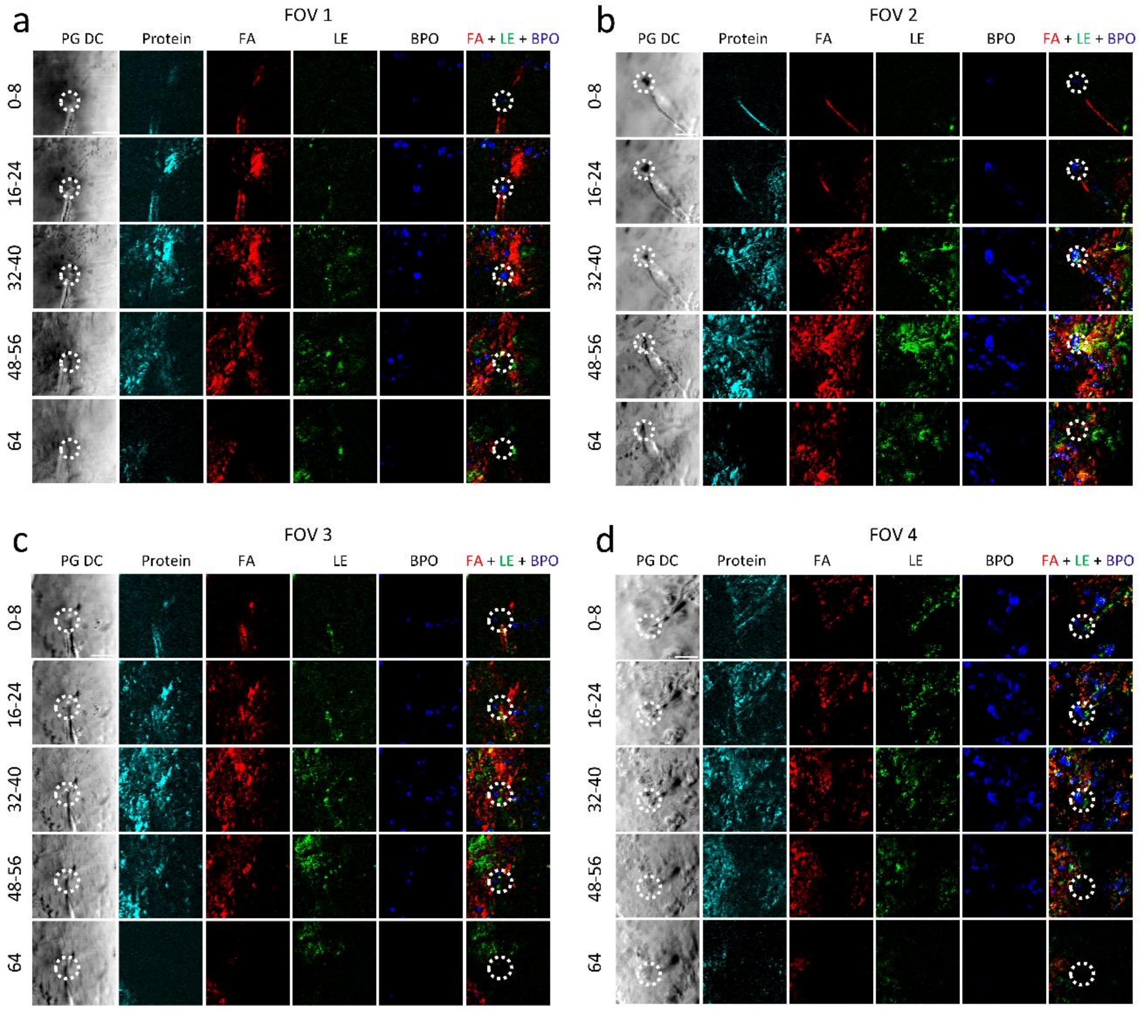


**Supplementary Figure 7. Depth-resolved phase gradient DC images and PTPG images at four field of views for unveiling drug pathway and quantitative evaluations of drug delivery efficiency**. Scale bar 50 µm. Field-of-view 180×180 µm$^2$. Probe power on sample was 10 mW. IR power on sample was ~2 mW with repetition rate 390 kHz, pulse width 100 ns. The imaging speed was 1.1 frame per second with a pixel dwell time of 10 µs. FA, fatty acid. LE, lipid ester. BPO, benzoyl peroxide.

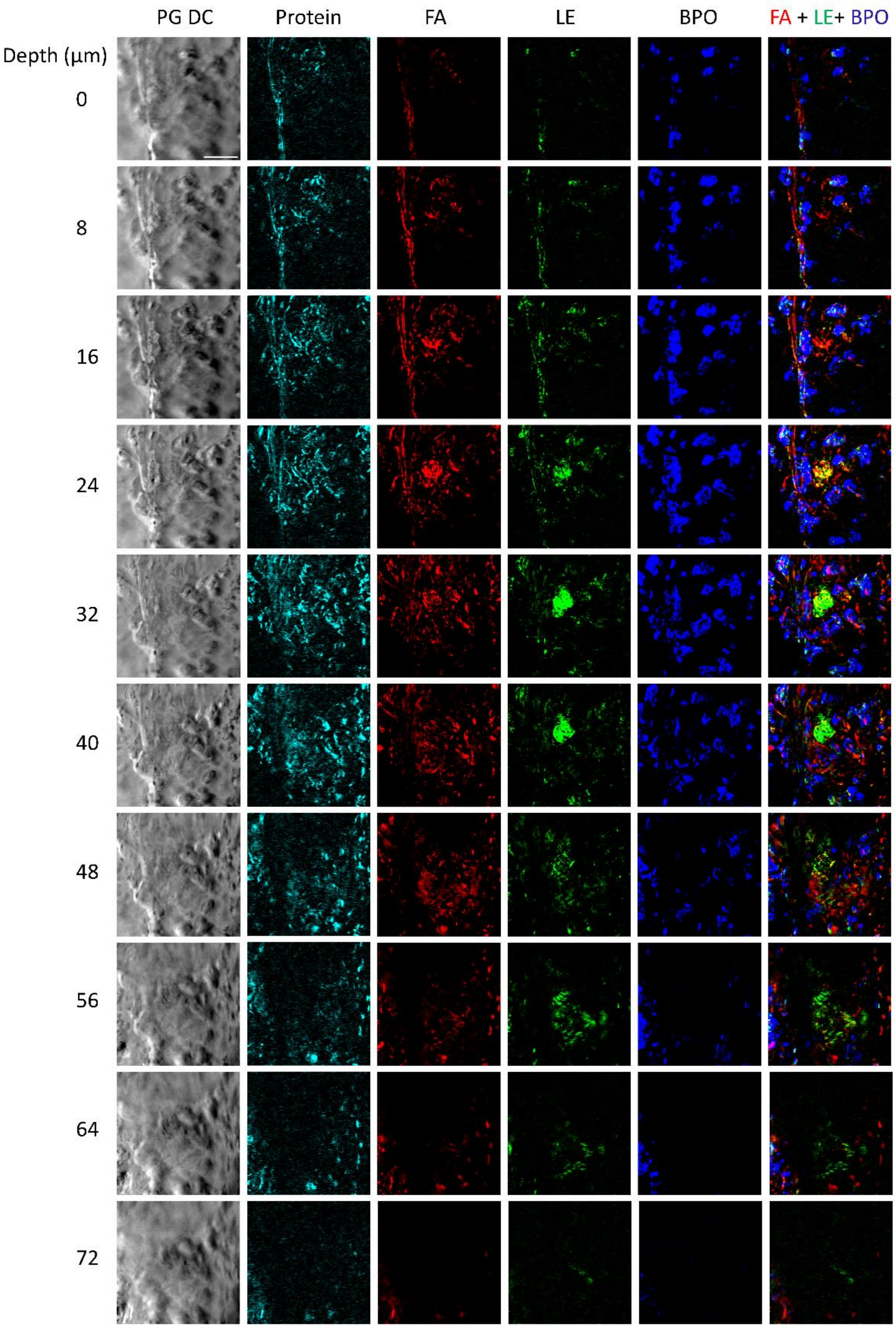


**Supplementary Figure 8. Depth-resolved images illustrating endogenous chemicals, including sebaceous glands, and the distribution of topical BPO beneath the skin**. The protein images were acquired at 1553 $cm^{-1}$, corresponding to the Amide II band absorption. FA, fatty acid. LE, lipid ester. BPO, benzoyl peroxide. Scale bar 50 µm.

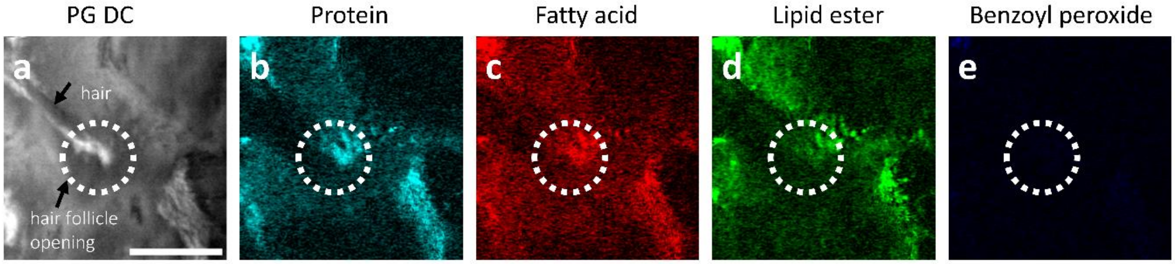


**Supplementary Figure 9. In vivo OPTM imaging of human skin without benzoyl peroxide administration**. **a**, phase gradient DC image of the stratum corneum showing a hair on the skin surface and the hair follicle opening. **b-e**, photothermal phase gradient images of protein (**b**, wavenumber 1553 $cm^{-1}$), fatty acid (**c**, 1704 $cm^{-1}$), lipid ester (**d**, 1742 $cm^{-1}$), and benzoyl peroxide (**e**, 1760 $cm^{-1}$).

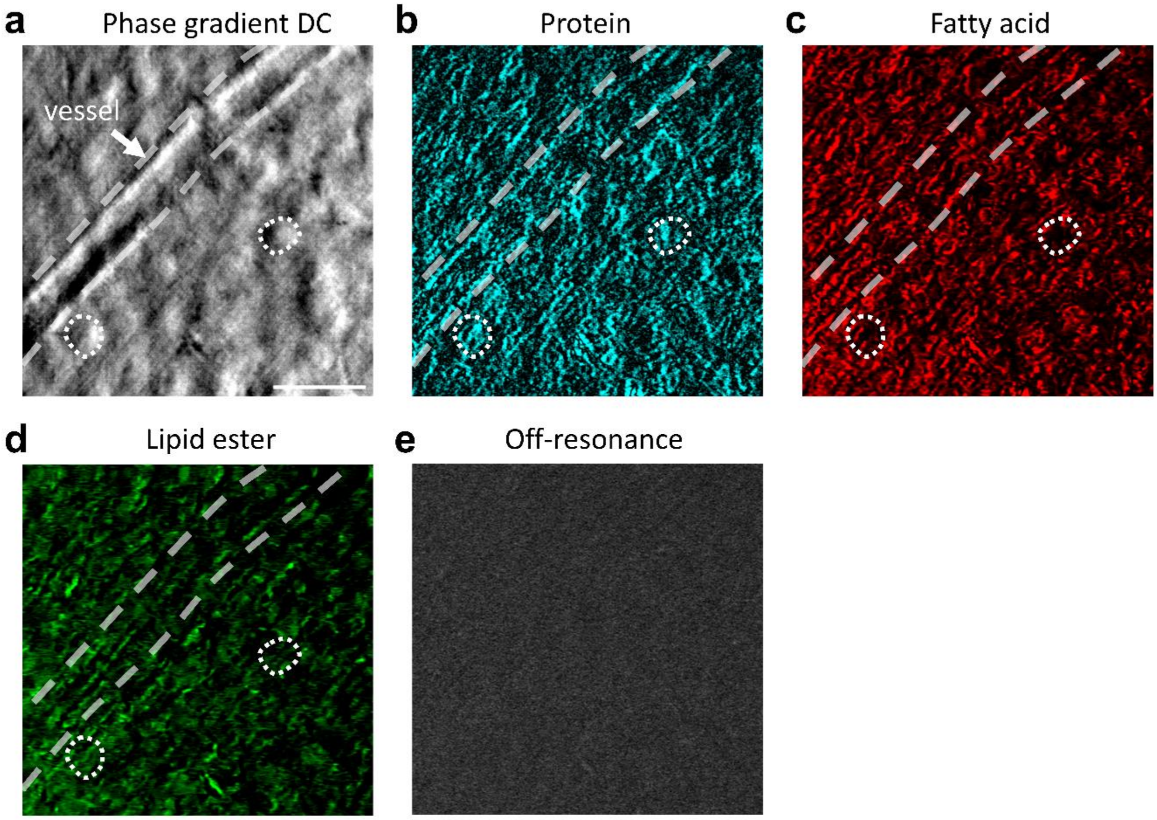


**Supplementary Figure 10. OPTM imaging of a mouse brain slice**. A 500-μm thick coronal brain slice was placed between a calcium fluoride ($CaF_2$) glass substrate and a coverslip, then positioned on a Teflon block to serve as a scattering substrate. Images were acquired by OPTM in an epi configuration. **a**, Phase gradient DC images of the brain slice. The blood vessel is outlined by dashed white lines, and two representative soma cells are marked with dashed white circles. **b-d**, Photothermal phase gradient (PTPG) images of endogenous chemical components. **b**, Protein image was acquired at IR wavenumber 1657 $cm^{-1}$, contributed by the absorption of Amid I band. The blood vessel wall is visualized to have rich protein content, which is contributed by elastin and collagen. **c-d**, Fatty acid and lipid ester images were acquired at 1704 and 1742 $cm^{-1}$, contributed by the absorption of acidic and esterified C=O bond in lipids. The endogenous protein and lipid contents within soma cells were detected. **e**, Off-resonance image was acquired at 1770 $cm^{-1}$ to validate the bond-selective imaging of OPTM. Scale bar 30 μm.

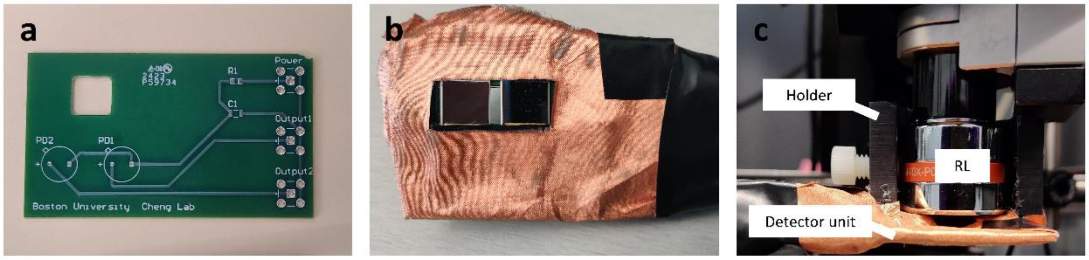


**Supplementary Figure 11**. **a**, a photo of the lab-built print circuit board (PCB). **b**, photo of split detector mounted on the PCB board. The detector unit was covered by copper mesh for isolating the external electrical noise. **c**, a photo of the detector unit mounted on the imaging objective via a 3D printed holder. RL: reflective objective.

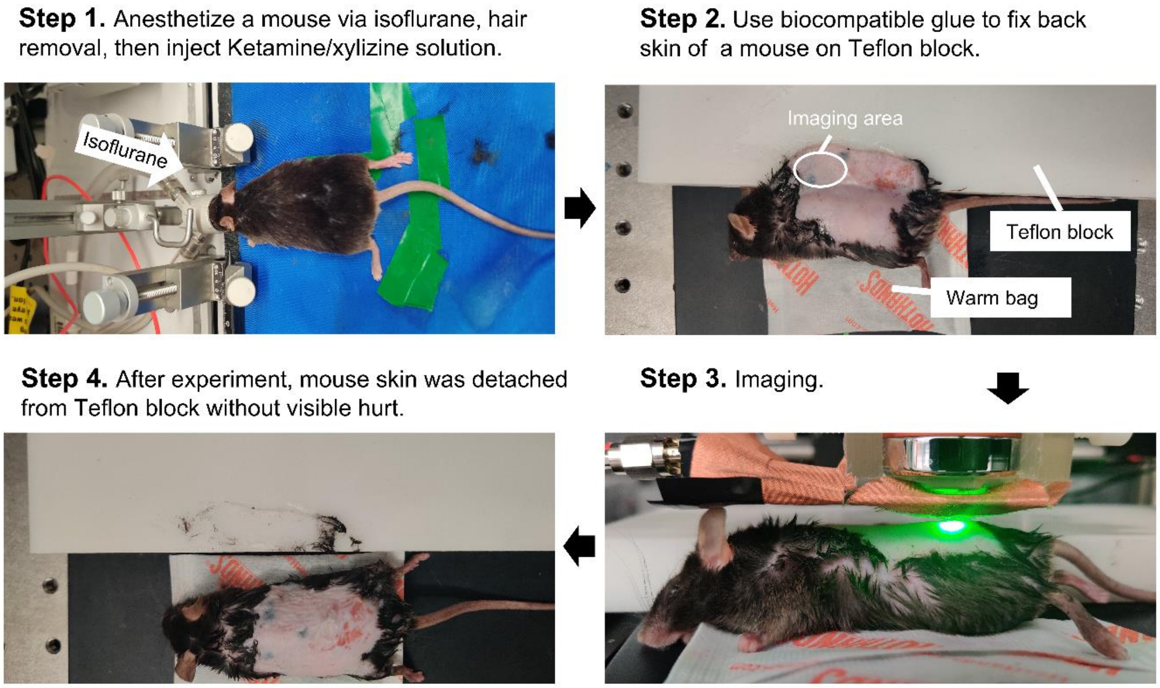


**Supplementary Figure 12. In vivo imaging of mouse skin.**

### Videos

**Movie S1.** Multispectral PTPG, absorption, and MIP images at a same region acquired from 1700 to 1770 $cm^{-1}$ with a step size of 2 $cm^{-1}$. Scale bar 30 µm. Field-of-view 150×150 $µm^2$.

**Movie S2.** 3D visualization of mouse ear skin after 5-µl drug sample administrated. This 3D view shows the hair and the hair follicles. Volume dimension 180*180*64 $µm^3$.

**Movie S3.** 3D visualization of mouse ear skin after 5-µl drug sample administrated. This 3D view includes a sebaceous gland under the skin surface. Volume dimension 180*180*72 $µm^3$.
